# A cell line model for the study of CD4-negative HIV-1 infection and latent virus reservoirs

**DOI:** 10.64898/2026.06.26.734695

**Authors:** Gizelle J. Lionel, Beth Binnington, Raymond W. Wong, Alan Cochrane, Jiazhen Jin, Donald R. Branch

## Abstract

Although controversial, limited publications support the notion that HIV-1 can infect CD4-negative cells. The objective of this study was to provide a comprehensive investigation of a universally available CD4-negative cell line model system that can be infected with X4 and R5 HIV-1 to generate integrated proviral DNA and serve to study latent viral reservoirs. The reason that HIV-1 infection of CD4-negative cells has become less investigated is due to a lack of a fully characterized model for the study of this unusual pathway. To address this critical need, human osteosarcoma (HOS) cells, engineered to express either CD4, CCR5 or CXCR4, and easily available from a commercial source were used. CD4 expression was examined using western immunoblot, flow cytometry, anti-CD4 blocking antibody and mRNA expression. Cells were infected with HIV-1 pseudo-enveloped viruses bearing either JR-FL (R5-tropic) or HXB2 (X4-tropic) envelopes, constructed on NL4-3 luciferase/GFP backbone. Infection was monitored by luciferase readout and visualized by GFP immunofluorescence. Raltegravir was used to inhibit integration, and AMD3100 and maraviroc used to block chemokine coreceptors, CXCR4 and CCR5, respectively. Productive versus latent infection was quantified by dual-fluorescence readouts using HI.fate.E. We confirmed that HOS cells lack CD4. HOS cells expressing only CCR5 or CXCR4 supported HIV-1 infection, although infection was significantly lower than in matched CD4-positive controls. Raltegravir treatment blocked proviral integration in all instances. Coreceptor antagonism and envelope-deficient viruses revealed that infection of CD4-negative CXCR4 cells remained CXCR4-dependent, whereas CD4-negative CCR5 cells showed evidence of CCR5-independent infection. Dual-reporter HI.fate.E assays indicated that CD4-negative cells could support both productive and latent infection. These studies establish a universally available cell line model for the study of CD4-negative HIV-1 infection. This cell line model will provide insight into the question of how CD4-negative cells can be infected with HIV-1 and whether CD4-negative cells can provide latent viral reservoirs in HIV/AIDS.

**Author summary:** Since the first description of HIV/AIDS in 1981 and the recognition that CD4 was a primary receptor for HIV-1 in 1983, a limited number of reports have suggested that cells lacking CD4 could be infected with HIV-1. These reports continued even when it was shown in 1996 that co-receptors, CXCR4 and CCR5, were also required for HIV-1 infection of CD4 T-helper cells. Indeed, crystallography studies showed that CD4 was required to interact with the HIV-1 envelope gp120 in order to cause conformational changes in the envelope to expose the binding motif for chemokine co-receptor engagement, required for additional conformational changes to expose the gp41 fusion protein, allowing for entry and infection. However, reports continued that cells lacking CD4 could be infected which raised questions as to how this can happen. To address this critical gap, we have identified a cell line, HOS, that is commercially available, having expression of CD4, CXCR4 and/or CCR5. Using these HOS cell lines, we have been able to confirm that HIV-1, either X4 or R5 enveloped viruses, can infect CD4-negative cells. We have also confirmed that infection is productive and allows for latent proviral integration. Our findings provide a system for further studies of the mechanism(s) of HIV-1 infection of CD4-negative cells using a consistent model and may aid in elucidating establishment of viral reservoirs.

## 1. Introduction

In the current dogma, HIV-1 entry into host cells is facilitated by the virus’ gp120 glycoprotein binding to the CD4 receptor on the cell surface [1,2]. This interaction induces a conformational change in the viral envelope, allowing gp120 to bind to one of the chemokine coreceptors, either CCR5 or CXCR4, found on susceptible cells like T cells and macrophages [2–7]. This interaction leads to additional conformational changes in gp120 that leads to exposure of gp41 (containing a fusion peptide) which facilitates fusion of the viral and cell membranes [2,7]. Although CD4 is the well characterized primary receptor for HIV, studies have shown that HIV can infect cells lacking detectable CD4 expression [8–24]. This evidence of CD4-independent infection pathways challenges the traditional understanding of viral entry mechanisms and suggest the use of alternative receptors or other factors that facilitate viral attachment and fusion. Several mechanisms have been proposed to explain how HIV 1 can infect CD4 negative cells across various targets. These include mutations in the gp120 envelope protein and direct engagement of coreceptors [9,10,11,12,13,14], use of alternative receptors such as galactoceramide [15,16,17], DC SIGN[18,19,20], or the mannose receptor[21,22], envelope independent pathways[23] or simply endocytosis [24]. Together, these findings suggest that HIV 1 can exploit multiple entry routes beyond the canonical CD4 mediated pathway. Whether CD4-negative cell infection is important for the pathophysiology of HIV-1 infection remains controversial.

Combination antiretroviral therapy (cART) effectively suppresses HIV-1 plasma levels to clinically undetectable level [25,26,27,28]. However, long-lived latent reservoirs within resting memory CD4^+^ T cells harbor replication competent viruses, representing a major obstacle to virus eradication [29, 30, 31]. Whether infection of CD4-negative cells presents another reservoir for latent HIV-1 provirus has not been fully addressed. In the study herein, we aim to identify a readily available cell line, human osteosarcoma cells (HOS), to serve as a model cell line to examine HIV-1 infection and latency. This cell line is easily obtained from commercial sources and engineered to express CD4 along with of the chemokine coreceptors, CXCR4 or CCR5, making it possible to study productive and latent viral infection and identify alternative routes that HIV-1 might use to infect epithelial cells that do not express CD4. The insights gained could have broader implications for understanding viral transmission, persistence in mucosal reservoirs, and challenges in completely suppressing HIV-1 infection.

## 2. Results

### 1. HIV-1 infection of CD4-positive and CD4-negative HOS cell lines and confirmation of lack of CD4

First, we assessed HIV-1 infectivity in CD4-negative HOS cells. We performed infection assays using luciferase reporter viruses matched to coreceptor usage (Fig. 1A). Based on luciferase readout, HOS-CD4 CCR5□ and HOS-CD4 CCR5□ cells were infected with the R5-tropic JRFL strain, while HOS-CD4 CXCR4□ and HOS-CD4 CXCR4□ cells were infected with the X4-tropic HXB2 strain. CD4-positive control cells exhibited robust infection, whereas CD4-negative cells were infectable but showed reduced infection efficiency compared to their CD4-positive counterparts. With R5, infection in CD4-negative cells was approximately two orders of magnitude lower than in CD4-positive cells, and with X4 it was approximately three orders of magnitude lower. Before we can conclude that CD4-negative cells were infectable, we needed to convincingly show that these cells were, in fact, devoid of CD4. We first performed a western immunoblot with anti-CD4. Figure 1B shows that HOS cells lack CD4 protein where various T-cell lines served as positive controls. Flow cytometric analysis confirmed distinct receptor expression profiles across our HOS cell panel (Table 1) (see Supplemental Materials, Figs. S1 and S2) for gating strategy and flow diagrams. HOS-CD4 CCR5□ and HOS-CD4 CXCR4 cells showed 66% and 32.6% CD4□ cells, respectively, alongside near-complete coreceptor expression (>95%). In contrast, HOS-CD4 CCR5□, HOS-CD4 CXCR4□, and wild-type HOS cells lacked detectable CD4 surface expression, while retaining high CXCR4 (endogenous) and/or CCR5 expression. The CD4-negative subpopulations in transduced lines were not due to insufficient antibody staining, as SupT1 T cells were 100% CD4□ and Raji B cells were CD4 (Fig. S2). It should be noted that wild-type HOS cells express endogenous CXCR4. Using an anti-CD4 blocking antibody, ibalizumab, also confirmed a lack of CD4 required for infection (Fig. 1C). Ibalizumab significantly reduced infection of HOS cells expressing CD4 but had no effect on infection of CD4 CCR5□ or CD4 CXCR4□ cells. Finally, we examined CD4 mRNA expression in the HOS cells. Figure 1D shows that HOS cells do not express CD4 mRNA. HOS-CD4 CXCR4□ and HOS-CD4 CCR5□ cells showed Cq values comparable to SupT1 controls (Cq = 27.2; positive threshold Cq < 38.2), while HOS-CD4 CXCR4□, HOS-CD4 CCR5□, and wild-type HOS cells had undetectable Cq values (negative threshold Cq > 38.2), matching Raji negative controls (Cq = 38.2). Taken together, these data establish a validated panel of CD4-negative HOS cells competent for HIV-1 infection, enabling dissection of CD4-independent entry mechanisms.

**Fig. 1.**
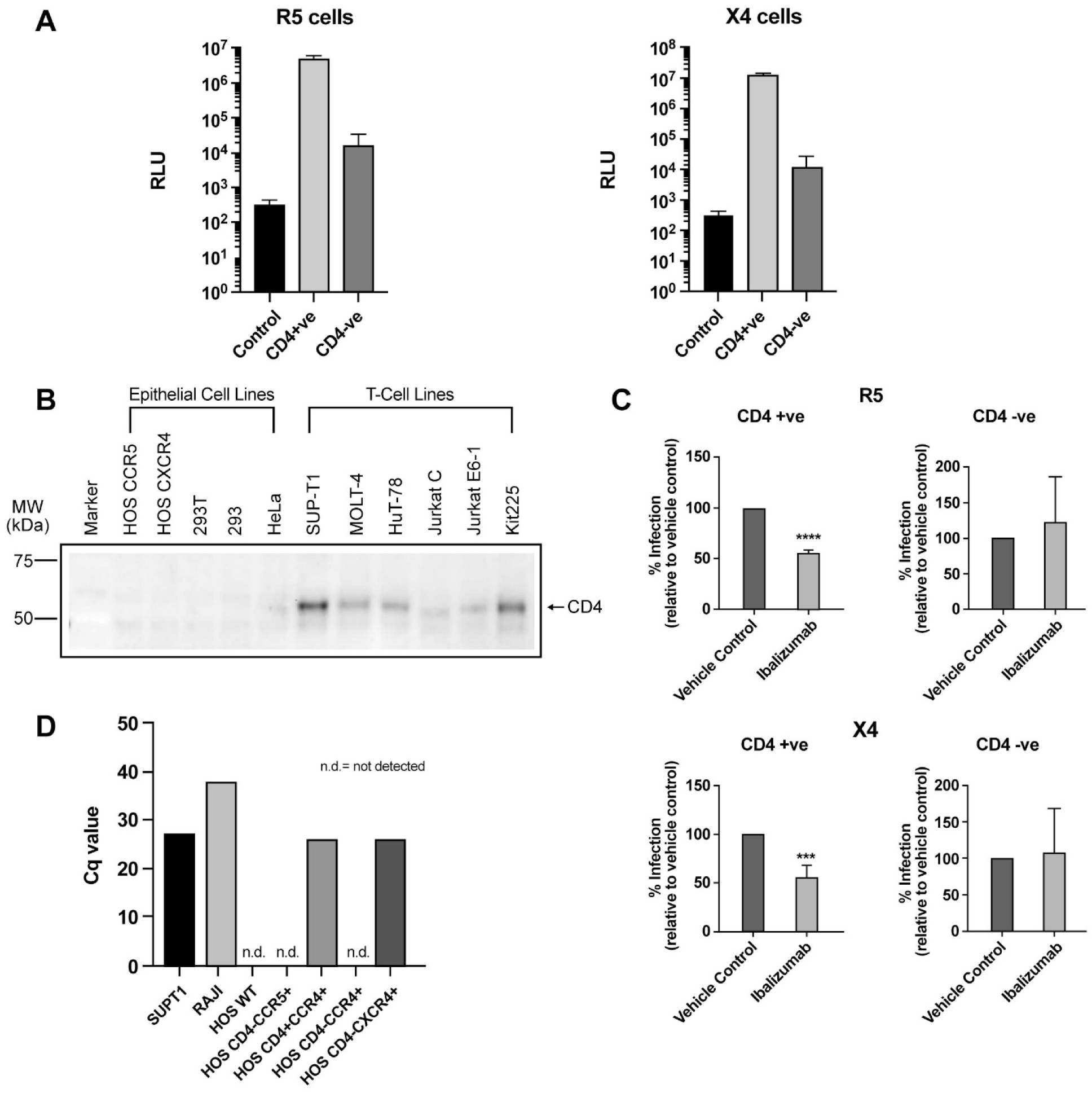
CD4-negative HOS cells support HIV-1 infection with confirmation of lack of CD4 protein and mRNA. **(A)** HOS cells expressing CCR5 (R5 tropic virus; left) HOS-CD4□CCR5□ and HOS-CD4□CCR5□ cells or CXCR4 (X4 tropic virus; right) HOS-CD4□CXCR4□ and HOS-CD4□CXCR4□ cells. Cells were infected with luciferase reporter virus containing JR-FL env or HXB2-env respectively, and infection was quantified by luciferase activity at 48 hours post infection. CD4-positive cells served as controls for infection. Luciferase reagent only (Control) is shown for each condition. Representative data from one of 3 independent experiments are shown. (**B**) Western blot for CD4 showing a lack of CD4 protein detected in HOS cells as well as other epithelial cell lines, including 293 and HeLa while T-cell lines served as controls for CD4 protein expression. (**C**) Anti-CD4 blocking antibody inhibits CD4^+^ but not CD4^−^ HOS cell infection. Cells were pretreated for 1 hour with 2 µg/mL Ibalizumab, with vehicle control (water control) included. RLU was measured 48 hours post-infection and expressed as percent infection relative to vehicle control. Experiments were repeated 3 times; a representative result is shown. (**D**) RT-qPCR quantification of CD4 mRNA. Cq values plotted (lower Cq = higher expression); SupT1 (Cq = 27.2) and Raji (Cq = 38.2) served as positive/negative controls. Cq > 38.2 considered negative; Cq < 38.2 considered positive.

**Table 1.** Summary of flow cytometry results for the cell-surface expression of CD4, CXCR4 and CCR5 on HOS cells.

| Cell Lines | CD4% | CCR5% | CXCR4% |
| --- | --- | --- | --- |
| HOS WILDTYPE | 0.0 | 97.826 | 10.926 |
| HOS CD4<sup>+</sup>CCR5<sup>+</sup> | 63.90-65.939 | 99.90 | 27.82 |
| HOS CD4<sup>-</sup>CCR5<sup>+</sup> | 0.0 | 99.966 | 48.516 |
| HOS CD4<sup>+</sup>CXCR4<sup>+</sup> | 32.316-32.6 | 32.0 | 100.0 |
| HOS CD4<sup>-</sup>CXCR4<sup>+</sup> | 0.0 | 2.454 | 99.4 |

### 2. Infection of CD4-negative HOS cells is low compared to CD4^+^ HOS cells

As seen in Figure 1A, the infection of HOS CD4^+^ cells appears to be much higher than infection of HOS CD4^−^ cells. To confirm this, we also examined infection of HOS CD4^+^CXCR4^+^ and HOS CD4^−^CXCR4^+^ using X4 virus expressing GFP. Figure 2 shows the results as measured by immunofluorescence and confirms that infection of CD4-negative cells is only about 5% of CD4-positive cells (quantitation confirmed via flow cytometry; see Supplemental Materials; Fig. S3).

**Fig. 2.**
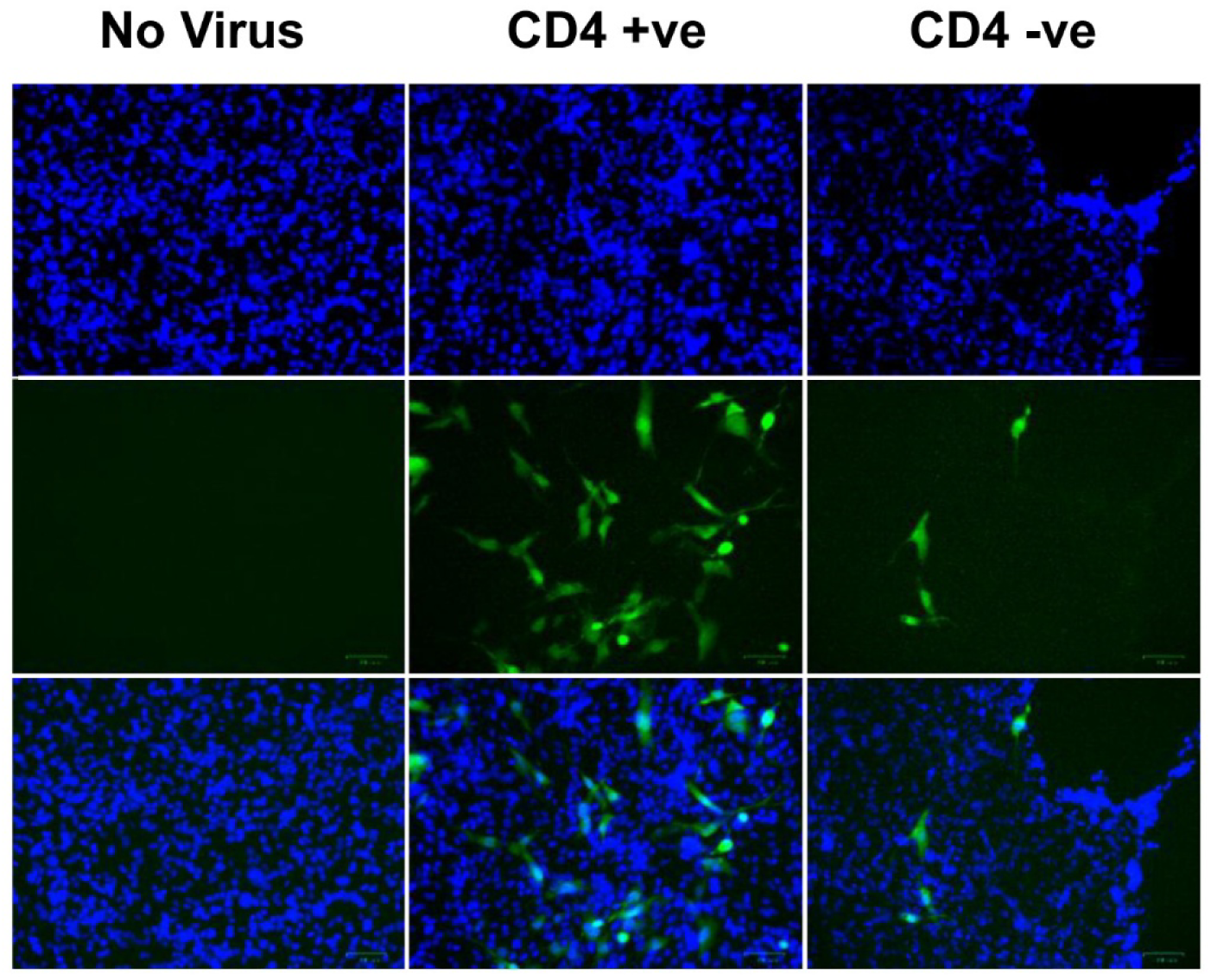
CD4^−^ HOS CXCR4^+^ cells show less infection than CD4^+^CXCR4^+^ HOS cells. Cells were infected with the HIV-1 GFP Reporter virus and imaged at 72 hours post-infection using fluorescence microscopy. A substantially higher population of GFP-positive cells was observed in the CD4-positive group compared to the CD4-negative cells. Notably, GFP expression in CD4-negative cells confirms viral integration, demonstrating that these cells support HIV-1 infection.

### 3. Raltegravir confirms HIV-1 integration in CD4-negative HOS cells

To confirm that HIV-1 infection of CD4-negative HOS cells reflects true proviral integration rather than non-specific reporter activity, we performed infection assays in the presence of the integrase inhibitor raltegravir (Fig. 3A). HOS-CD4□ CXCR4□ and HOS-CD4 CXCR4□ cells were infected with luciferase reporter virus ± raltegravir. In both CD4□ and CD4□ cells, raltegravir completely inhibited luciferase expression, demonstrating that the observed luciferase activity requires HIV integrase-mediated strand transfer and integration into the host genome. These findings validate that CD4-negative HOS cells support genuine HIV-1 integration, ruling out possible non-integrative artifacts.

**Fig. 3.**
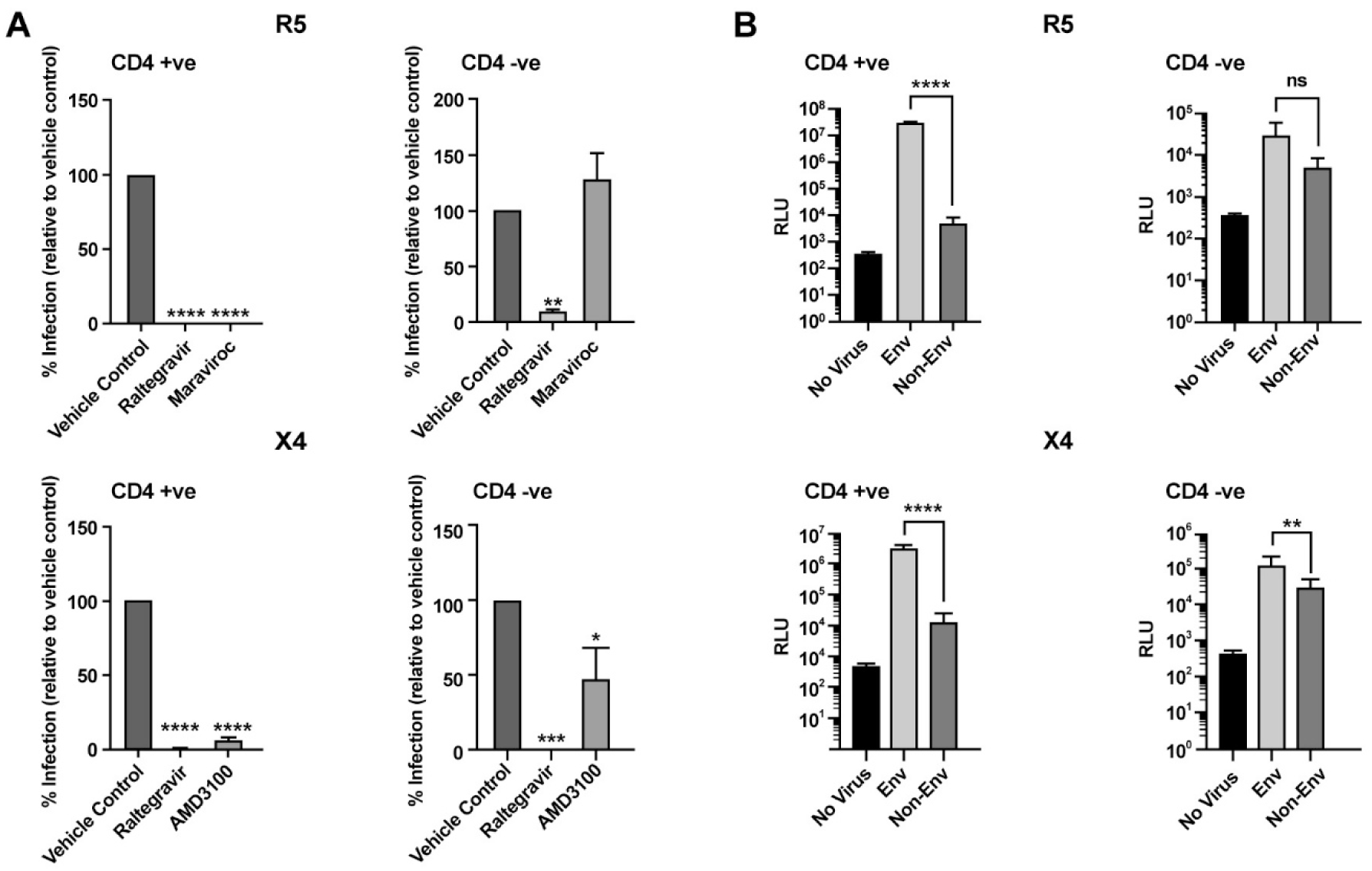
HIV-1 infection of CD4-negative HOS cells shows differential coreceptor and envelope dependence. HOS cells were infected with luciferase reporter viruses in the presence of coreceptor antagonists or with virus-not containing envelope. (A)Top: Infection of HOS-CCR5 cells with Maraviroc and Raltegravir & Bottom: Infection of HOS-CXCR4□ cells with Plerixafor (AMD3100) and Raltegravir. Cells were pretreated for 1 hour with either 1 µM Raltegravir, 1 µM Maraviroc or 1 µM AMD3100 with vehicle control (0.1% DMSO) included. RLU was measured 48 hours post-infection and expressed as percent infection relative to vehicle controls. Vehicle controls yielded readouts of 3.2 × 10^7^ RLU (CD4-positive) and 3.7 × 10^4^ RLU (CD4-negative) in Top, and 4.2 × 10^6^ RLU (CD4-positive) and 9.0 × 10^4^ RLU (CD4-negative) in Bottom. In CD4-positive cells, Raltegravir, Maraviroc, and AMD3100 all produced significant inhibition. Raltegravir achieved near-complete inhibition in CD4-negative cells, and AMD3100 reduced infection in CD4-negative X4 cells by ∼50%. Experiments with each drug were repeated 3 times; a representative result is shown. **(B)** HOS-CD4□CCR5□ and HOS-CD4□CCR5□ cells (Top) and HOS-CD4□CXCR4□ and HOS-CD4 CXCR4□ cells (Bottom). Cells were infected with HIV-1 luciferase reporter virus containing HIV-1 envelope (Env) or lacking envelope (Non-Env). 48 hours post infection RLU was measured. “No virus” wells (cells plus reagents, no virus) served as negative controls. Data are presented as luciferase activity. Significantly higher infection was observed in CD4-positive cells exposed to envelope-containing virus compared to envelope-deficient controls. In CD4-negative R5 cells (Top right), there was no significant difference in the level of infection with or without the envelope added. CD4-negative X4 cells (Bottom right) indicated a significantly higher level of infection with the addition of envelope demonstrating that HIV-1 infection in these CD4-negative cells is envelope-dependent.

### 4. Specific chemokine co-receptor usage

To determine whether HIV-1 infection of CD4-negative HOS cells depends on canonical coreceptor usage, we treated cells with the CXCR4 antagonist plerixafor (AMD3100) and the CCR5 antagonist maraviroc during infection. Figure 3A shows that for HOS-CD4 CXCR4 cells, plerixafor reduced infection, indicating that HIV-1 entry in this context remains CXCR4-dependent despite the absence of CD4. In contrast, maraviroc had no significant effect on infection of HOS-CD4 CCR5□ cells, suggesting that CCR5 is not required for HIV-1 entry into these cells under our experimental conditions.

### 5. Envelope dependence of HIV-1 infection in CD4-negative HOS cells

Results from Figure 3A showing a lack of CCR5-dependent infection by R5 virus led us to examine whether infection of CD4-negative cells required the viral envelope glycoprotein. We used cells infected with luciferase reporter viruses carrying either HIV-1 Env (Env) or lacking Env(Non-Env “core” virus), using the same infection protocol as above, with “no virus” wells serving as negative controls (Fig. 3B). In HOS-CD4 CCR5□ and HOS-CD4 CXCR4□ cells, envelope-containing virus yielded infection levels approximately 10³-fold higher than non-Env virus, confirming robust envelope-dependent entry in CD4□ controls. In HOS-CD4 CXCR4 cells, Env virus also produced significantly higher infection than non-Env virus, indicating that infection of these CD4-negative cells is largely envelope-dependent. By contrast, in HOS-CD4 CCR5□ cells, non-Env virus did not show significantly reduced infectivity compared to Env virus, consistent with an envelope-independent infection route in this population. Together, these findings suggest that HIV-1 can exploit a CD4-independent, CCR5-independent, and envelope-independent pathway in HOS-CD4 CCR5□ cells, while infection of HOS-CD4 CXCR4□ cells remain dependent on CXCR4 and the viral envelope.

### 6. CD4-negative HOS cells support productive and latent HIV-1 infection

To assess whether HIV-1 infection of CD4-negative HOS cells can give rise to productive and latent infection, we used the HI.fate.E reporter virus pseudotyped with HXB2 Env to infect CD4 CXCR4 and CD4 CXCR4□ HOS cells. HI.fate.E expresses E2 Crimson from the HIV-1 LTR and ZsGreen from an internal constitutive promoter, enabling discrimination of uninfected (double-negative), productively infected (E2 Crimson /ZsGreen^+^), and latently infected (ZsGreen□ only) cells by flow cytometry [38]. Seventy-two hours post-infection, approximately 2–3 × 10□ cells were harvested per condition, and 5 × 10□ events were acquired per sample to capture rare infection events, particularly in the CD4-negative population.

Consistent with previous characterization of this reporter, cells in the P5 gate (Q2+Q3; Crimson□ with or without ZsGreen) were interpreted as productively infected, whereas cells in the P6 gate (Q4; ZsGreen□ only) were classified as latently infected. In Figure 4A, HOS-CD4 CXCR4□ cells, 2.3% of the population fell within P5 and 0.3% within P6, indicating that a subset of infected cells enter latency following productive infection. In Figure 4B, HOS-CD4 CXCR4□ cells, infection frequencies were markedly lower, with 0.1% of cells in P5 and 0.02% in P6, but the same pattern of a minority of infected cells adopting a latent phenotype was observed. These preliminary data suggest that once HIV-1 gains entry, the likelihood to establish latency relative to productive infection is qualitatively similar in CD4-positive and CD4-negative HOS cells, even though overall infection efficiency is substantially reduced in the absence of CD4.

**Fig.4.**
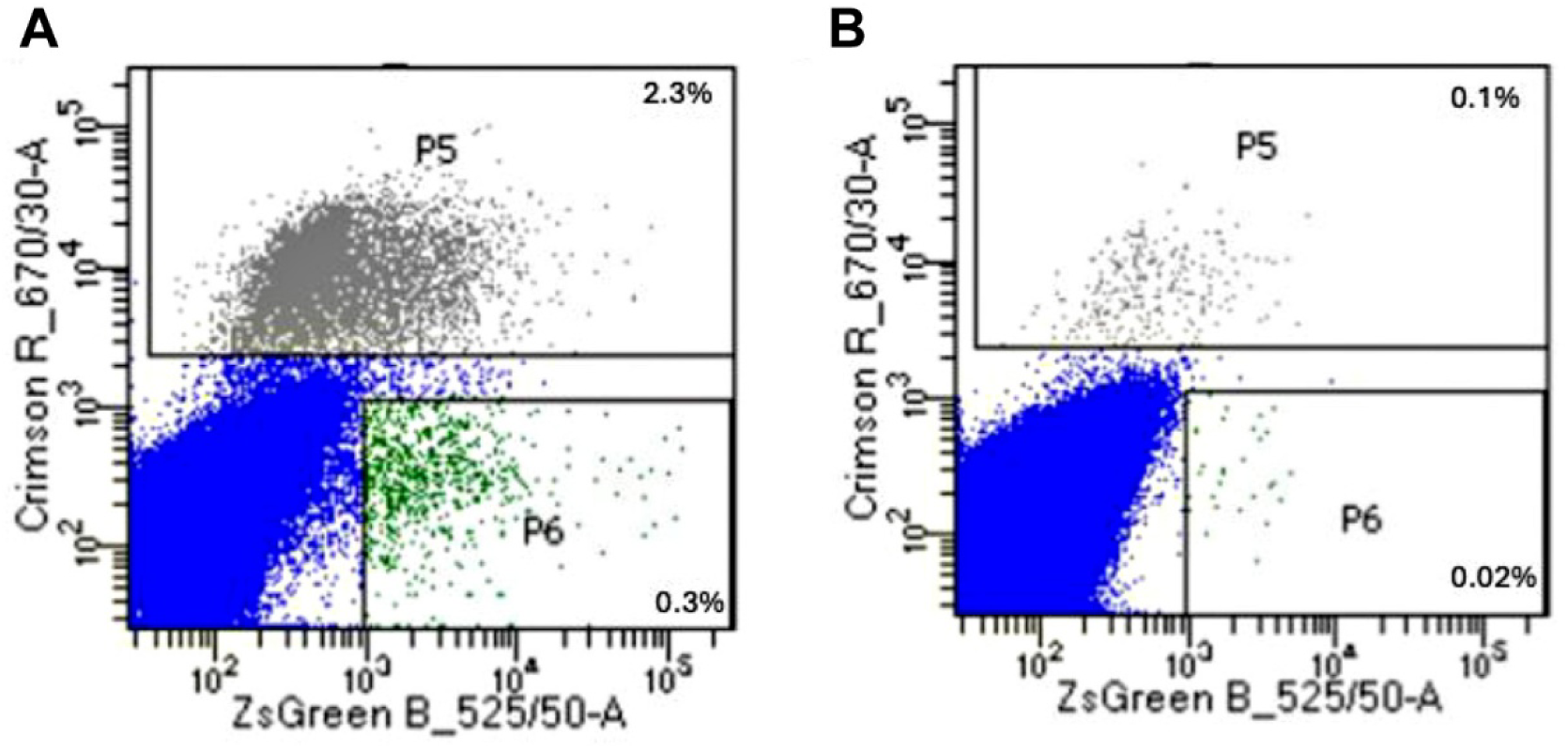
HI.fate.E reporter analysis of productive and latent HIV-1 infection in CD4-positive and CD4-negative HOS cells. HOS-CD4□CXCR4□ and HOS-CD4□CXCR4□ cells were infected with HXB2-pseudotyped HI.fate.E virus.Cells were acquired for flow cytometry analysis 72hours post infection. Representative two-parameter plots of E2 Crimson versus ZsGreen fluorescence. Quadrants were defined as follows: Q1 (double-negative) uninfected cells; P5 (Q2+Q3; E2 Crimson with or without ZsGreen) productively infected cells; P6 (Q4; ZsGreen only) latently infected cells. (A) In CD4□CXCR4□ cells, 2.3% of cells were productively infected (P5) and 0.3% were latently infected (P6). (B) In CD4□CXCR4□ cells, 0.1% of cells were productively infected and 0.02% were latently infected. Data shown are presented as percentages of total live, single cells.

## 3. Discussion

The objective of this work was to demonstrate that CD4-negative cells are permissive to HIV 1 infection and provide a cell line model for further studies. HOS cells were selected over other potential cell lines due to the universal availability of these cells expressing CD4, CXCR4 or CCR5 in various combinations. We confirmed that HOS cells completely lack CD4 expression, including at the mRNA level, and that the cells are productively infectable by HIV-1, both X4 and R5 viruses, as indicated by luciferase production and inhibition of luciferase expression using raltegravir. Using HI-fateE virus, we provide evidence that CD4-negative HOS cells can be latently infected. Thus, we have achieved our objective in describing a universally available cell line that is CD4-negative which can be productively infected with X4 or R5 HIV-1. Infection of CD4□ negative cells was completely inhibited by raltegravir, indicating that entry and subsequent luciferase or GFP reporter expression reflect integrase□ dependent proviral integration rather than non□ specific reporter activity. These observations add to growing evidence that HIV□ 1 can infect CD4□ deficient targets under certain conditions and reinforce the concept that CD4 is not absolutely required once alternative entry routes are available [23,42,43].

Our coreceptor and envelope involvement experiments revealed different entry requirements between CD4 negative CXCR4□ and CCR5□ HOS cells. In CD4^−^CXCR4□ cells, infection remained sensitive to CXCR4 antagonism and was markedly reduced with envelope□ deficient virus, supporting an entry pathway that is CD4□ independent but still CXCR4□ and Env dependent [9]. In contrast, infection of CD4^−^CCR5□ cells were not significantly affected by CCR5 blocker maraviroc and showed comparable levels with Env□ containing and Env□ deficient virions, consistent with a CD4 independent, CCR5 independent, and envelope independent mode of entry. This suggests that distinct cellular pathways or alternative receptors may facilitate HIV□ 1 uptake in different CD4 negative contexts, and that CCR5 expression in these cells is not the primary determinant of permissiveness.

Previous studies have demonstrated an HIV-1 envelope-independent mode of entry and infection in epithelial cells and other CD4-negative cells [23,42,43] like observations in our study. A study looking at the CD4-independent infection of trophoblasts used the NL4-3-Luc E-R+ expression vector to generate Env-deficient reporter viruses, and the results showed HIV-1 infection of these cells independent of envelope [42]. In the study by Pang et al. [23], gingival epithelial cells(normal human oral keratinocytes) were infected with HIV-1 NL4-3-EGFP Env^−^(no envelope added) and HIV-1 NL4-3-EGFP Env^−^with HIV-1 LAI envelope proteins. Their results indicated that similar levels of infection were observed with or without the addition of envelope in the CD4-negative cells, but CD4^+^ HeLa cells were not infectable with the HIV-1 NL4-3-EGFP Env^−^ only, an indication that envelope-independent infection occurred only in CD4-negative cells [23]. The study included the use of other cell types including a prostate cell line, and an epithelial-like fibrosarcoma cell line, like the HOS cells used in this study. There was a significant level of infection of these CD4-negative cells with the virus lacking an envelope, although the efficiency of this HIV-1 infection was low [23]. Therefore, the findings of our study align with previously reported results indicating that HIV-1 can infect CD4-negative cells via an envelope independent mechanism. In the study by Pang et al. there are no direct investigations to prove that the cells were CD4-negative, except infection using HIV-1 gp120 monoclonal antibody, which did not inhibit infection in the CD4-negative cells. However, in our study, we demonstrate that this mode of infectivity happens without any CD4 involvement. It is important to emphasize that the Env independent productive infection observed in our CD4 negative cells is not unique to this context; similar Env independent or partially Env independent entry has been reported in other cell systems [42,43], and recent work further demonstrates that HIV 1 can fuse within pH neutral endocytic vesicles in both cell lines and primary CD4^+^ T cells [44,45,46]. Although we could not exclude endocytosis as a potential entry mechanism within this study period, future experiments will specifically investigate this pathway.

Using the HI.fate.E dual reporter system, we further show that HOS-CD4^−^CXCR4 cells not only support productive infection but also establish a detectable latent reservoir, albeit at substantially lower absolute frequencies than CD4 positive cells. The relative proportion of infected cells that adopt a latent phenotype was broadly similar between CD4 positive and CD4 negative populations, suggesting that once integration has occurred, the decision between productive replication and latency may be governed by host and viral factors that are largely independent of CD4 expression at entry. These preliminary data raise the possibility that non canonical target cells could contribute to latent infection under certain in vivo conditions, even if they are infected inefficiently [47].

In our experiments, both CD4-positive control and CD4-negative cells showed low levels of infection, a pattern consistent with clinical observations that only a small proportion of CD4^+^ T cells harbor HIV-1 in infected individuals [48], as well as in CD4□ independent infection of CD8+ lymphocytes [8]. Taken together, these studies indicate that low infectivity does not rule out genuine CD4-independent entry in our HOS CD4-negative cells but rather reflects the intrinsically low frequency of productive HIV 1 infection in many cell types.

In conclusion, we report a well characterized cell line model system that is easily obtainable from commercial sources, for the systematic study of HIV-1productive and latent infection of CD4-negative cells.

## 4. Materials and methods

### Cells

In these experiments, we used human osteosarcoma (HOS) cell lines, we received HOS wild type (CRL-1543) from ATCC. The following HOS cell lines were obtained through BEI Resources, NIAID, NIH (formally the NIH AIDS Reagent and Reference Reagent Program): HOS-CD4/CXCR4, HOS-CXCR4, HOS-CD4/CCR5, HOS-CCR5 from Dr. Nathaniel Landau (Cat# 3319,3518,3318,3519). HOS cells had been transduced with a retroviral vector encoding either coreceptor leading to the HOS-CCR5 and HOS-CXCR4 cells to stably express these co-receptors without CD4 expression. Additionally, HOS cells were stably transfected to express CD4, then these HOS CD4+ cells were transduced with a retroviral vector encoding either coreceptor, resulting in HOS cells co-expressing CD4 and CCR5 or CXCR4. These cell lines were cultured in Dulbecco’s Modified Eagle Medium (DMEM,Wisent, Saint-Jean-Baptiste, Canada), supplemented with 10% heat-inactivated Fetal Bovine Serum (FBS; Wisent, Saint-Jean-Baptiste, Canada), and 100 units/mL penicillin (Gibco, Burlington, Canada), 100 μg/mL streptomycin (Gibco). Human embryonic kidney cells (HEK 293T, ATCC, CRL-11268) were maintained in Dulbecco’s Modified Eagle Medium (DMEM; Wisent), supplemented with the same concentrations of FBS and antibiotics as described above. SUP-T1 [Smith et al. 1984] (a human T-lymphoblastic lymphoma cell line) and Raji [Pulvertaft et al. 1964] (a human B-cell lymphoma cell line) all from ATCC were cultured in complete medium consisting of RPMI 1640 1X with L-Glutamine & Sodium Bicarbonate (Wisent), supplemented with 10% fetal bovine serum (FBS), 2 mM L-glutamine, 25 mM HEPES, and 1% penicillin-streptomycin (P/S). All cells were incubated at 37°C in a 5% CO2, atmosphere.

### Plasmids

We were kindly gifted the HIV-1 NL4-3 ΔEnv Vpr Luciferase Reporter Vector (pNL4-3.Luc.R-E-) from Dr. Vineet KewalRamani, New York Medical University, NY. The firefly luciferase gene was integrated into the nef region of the pNL4-3 plasmid. Two frameshift mutations (one in the 5’ env region and one at amino acid 26 of Vpr) render this clone deficient in Env and Vpr proteins. This construct supports only a single cycle of viral replication. Dr. Vineet KewalRamani also gifted us the HIV-1 JR-FL Env and the amphotropic envelope glycoprotein of vesicular stomatitis virus (VSV-G) Env expression vectors. We were generously provided the HIV-1 HXB2 Env expressing vector from Dr. M. Tremblay, Quebec City, QC. The following reagent was obtained through BEI Resources, NIAID, NIH: pNL-GFPΔEnv Vector, HRP-20247.The plasmid pNL-GFPΔEnv was created by replacing the NLuc gene in NL-NLucΔEnv with the GFP reporter gene. We also received the HI.fate.E dual reporter plasmid as a generous gift from Dr. Thomas Murooka (University of Manitoba). The HI.fate vectors are derived from the HIV-1 pNL4-3.HSA.R E plasmid backbone. The HI.fate.E reporter vector contains two fluorescent markers: E2-Crimson, which is expressed from the HIV-1 long terminal repeat (LTR) and acts as an indicator of HIV-1 gene transcription, and ZsGreen, which is expressed from an internal promoter and marks cells where HIV-1 has entered and integrated into its genome. Plasmid sequences were verified using Plasmidsaurus (plasmidsaurus.com), a commercial nanopore sequencing service providing full-length plasmid sequencing.

### Reagents

The HIV-1 inhibitors included Raltegravir, Maraviroc, and Plerixafor (AMD-3100), all obtained from BEI Resources, and anti-CD4 blocking antibody, Ibalizumab (IgG4SP isotype) was purchased from NovoProLabs (Cat. #176843, Shanghai, China).

### Viruses

Recombinant, replication-incompetent pseudotyped HIV-1 particles bearing the luciferase reporter gene were generated using HXB2 (X4), JR-FL (R5), or VSV-G envelopes, as previously described.[Bokaei et al. 2007] Briefly, 8.5 × 10 HEK 293T cells were plated in 10 mL of DMEM media 24 hours prior to transfection. Co-transfection was performed by introducing 7.2 µg of HIV-1 NL4-3luc or pNL-GFPΔEnv or HI.fate.E together with 10.8 µg of plasmids encoding HXB2, JR-FL, or VSV-G envelope proteins into each 10 cm dish of HEK 293T cells. Transfections were performed using Lipofectamine® 3000 Transfection Reagent (Invitrogen, Thermo Fisher Scientific, Cat. No. L3000-015, Waltham, MA, USA), following the manufacturer’s instructions. DMEM media was changed after 6 hours, and on days 2 and 3 the viral supernatant was collected. The supernatant was filtered through sterile 0.45 µm Acrodisc® syringe filters with Supor® membrane (Pall Corporation, Port Washington, NY, USA) and the virus containing-supernatant was placed in the −80 in 1mL aliquots. Viral supernatants were used for most infection assays. For certain virus preparations, however, supernatants were further concentrated following filtration by underlaying with a 20% sucrose cushion and centrifuging at 16,000 × g for 1.5 hours at 4 °C. Resulting viral pellets were resuspended in TNE buffer (20 mM Tris-HCl, pH 7.5; 1 mM EDTA; 100 mM NaCl) and stored in aliquots at –80 °C. HIV-1 p24 capsid protein levels were quantified using an ELISA kit (ZeptoMetrix, REF0801002, Buffalo, NY). The p24 concentration was used to estimate the number of viral particles and to calculate the multiplicity of infection (MOI), based on the ratio of viral particles to target cells. Each virus preparation was subjected to a single freeze-thaw cycle prior to their use in infection experiments.

### HIV-1 infection

For HIV-1 infection assays, 30,000 cells were plated in a 96-well plate in 100 µL of media. There were 6 replicates per cell line. After the cells adhered, 100 µL of virus was added to all wells except one row of well dedicated to the negative control or background. Unless otherwise stated, typically CCR5+ cells were infected with JR-FL (R5 virus) and CXCR4+ cells were infected with HXB2 (X4 virus). At 48–72 hours post-infection, luciferase reagent (Promega, Cat# E4030, Madison, WI, USA) was added according to the manufacturer’s protocol, and luminescence was measured using a PerkinElmer plate reader, measured in relative light units (RLU).

For inhibition assays, the protocol was the same as above, except that cells were pretreated with the inhibitors for 1 hour prior to viral infection. 1mM drug stocks were prepared in DMSO or water and stored frozen at −30□. Final concentrations are as shown in figure legends. A vehicle control consisting of 0.1% DMSO or water was included in these infection assays. To visualize the percent of CD4-negative cells becoming infected, cells were infected with the pNL-GFPΔEnv-HXB2 virus. We followed this protocol: 200,000 cells were collected by centrifugation and resuspended in 500 µL of lentiviral transfection supernatant (409 ng p24/µL). The cell suspension was incubated at room temperature for 30 minutes to enhance viral binding, before being transferred to a 6 well plate containing 2mL of complete media. 72 hours post-infection, GFP-positive cells were observed under a fluorescence microscope (ZOE Fluorescent Cell Imager) and representative images were captured for analysis.

HI.Fate.E plasmid pseudotyped with HXB2 Env was used to infect HOS□ CXCR4 cells. A total of 5 × 10^5 cells were seeded per well in a 6 well plate, and 1 mL of viral supernatant was added to each well and incubated for 72 hours.

### CD4 gene expression

For qRT-PCR quantification, total RNA was isolated from approximately 1□×□10□ cells using the Aurum Total RNA Mini Kit (Bio-Rad, Cat. #732-6820, Hercules, CA, USA). cDNA synthesis was performed using the iScript cDNA Synthesis Kit (Bio-Rad, Cat. #170-8841) according to the manufacturer’s instructions. The resulting cDNA was diluted 2- to 10-fold in nuclease-free water prior to qPCR.

To detect CD4 expression, qRT-PCR was performed using CD4-specific primers previously described in Schenkel et al. 2014,[Schenkel et al. 2010] with the following sequences: CD4 Forward (5’GCCCTTGAAGCGAAAACAG3’), CD4 Reverse (5’CTCCTTGTTCTCCAGTTTCAAAC3’). For normalization, human ACTB (β-actin) expression was quantified using a validated primer set (OriGene Technologies, Cat. #HP204660, Rockville, MD, USA): hACTB Forward (5’–CACCATTGGCAATGAGCGGTTC–3’) and hACTB Reverse (5’–AGGTCTTTGCGGATGTCCACGT–3’).

qRT-PCR reactions were assembled in 20□μL volumes using SsoAdvanced Universal SYBR Green Supermix (Bio-Rad, Cat. #1725271), containing 5□μL of diluted cDNA, 300□nM of each primer, and nuclease-free water. Reactions were performed in a 96-well PCR plate (Bio-Rad, Cat. #HSP9601) sealed with adhesive qPCR seal (Sarstedt, REF 95.1999, Germany).

Amplification and fluorescence detection were carried out using the CFX96 Touch Real-Time PCR System under the following thermal cycling conditions: 95□°C for 30□s, followed by 45 cycles of 95□°C for 5□s and 60□°C for 15□s. Amplification data were analyzed using CFX Maestro software (Bio-Rad), and PCR products were further verified by agarose gel electrophoresis.

### Western blot

Western blot for CD4 was done using a panel of epithelial cells, including HOS, 293 and HeLa and T-cells, including SupT-1, Molt4, HuT-78, Jurkat C, Jurkat E6-1 and Kit225. Lysates were prepared from 10^7^ cells using radioimmunoprecipitation assay (RIPA) lysis buffer and loaded onto a 10% SDS-PAGE gel. The gel was transferred to nitrocellulose membrane and probed with polyclonal goat anti-human CD4 primary antibody (Novus Biologicals, R&D Systems #AF-379-NA, Lot #YS0819111) and protein visualized using rabbit anti-goat HRP (Novus Biologicals) and enhanced chemiluminescence.

### Flow Cytometry and data acquisition

Cell surface expression of CD4, CXCR4, and CCR5 was assessed by flow cytometry using 300,000 HOS cells per FACS tube. Surface staining was performed as previously described. Briefly, cells were incubated in 100 µL of staining buffer (1XPBS, 2% FBS, 1mM EDTA) containing 5 µL each of FITCconjugated mouse antihuman CD4 (BioLegend, Cat. #300514, San Diego, CA, USA), APCconjugated mouse antihuman CXCR4 (BioLegend, Cat. #539), and PEconjugated mouse antihuman CCR5 (BioLegend, Cat. #2614).

Data from 5000 events per tube were acquired using a CytoFLEX LX flow cytometer (Beckman Coulter, Loaner CytoFLEX LN, S/N BB21022, ID#81262408). Fluorescence signals were collected in the following channels: FITC (B525), PE (Y585), and APC (R660), along with forward scatter (FSC) and side scatter (SSC).

Flow cytometry data were analyzed using FlowJo v10.9.0. The gating strategy (illustrated in the supplemental figures; S1) was as follows: (1) side scatter area (SSC-A) versus forward scatter area (FSC-A) plots were used to select the primary cell population based on size and granularity and excluding debris; (2) forward scatter height (FSC-H) versus forward scatter area (FSC-A) gating was used to exclude doublets and aggregates; (3) side scatter height (SSC-H) versus side scatter width (SSC-Width) gating provided an additional doublet exclusion step for selection of single cells; (4) viability gating was performed using a live/dead dye to exclude dead cells, ensuring analysis was limited to live, single cells. Subsequent dual-parameter plots of CD4 versus CCR5 and CD4 versus CXCR4 confirmed receptor expression profiles for each HOS cell line.

For flow cytometric analysis usingHIV-1 HI.fate and GFP-reporter viruses (containing the HXB2 envelope) we infected X4 cells, approximately 2–3 million cells were collected 72 hours post-infection. The HI.fate virus expresses both Crimson Red and ZsGreen fluorescent proteins, and flow cytometry was performed using BD LSRFortessaTM X-20 II at the University of Toronto Centre for Immune Analytics Core Facility, collecting 250,000-400,000 events. Dual-parameter plots of Crimson versus ZsGreen fluorescence were used to distinguish cells with active viral gene expression from those harboring integrated but latent virus. Data analysis followed the gating strategy described above.

### Statistical analysis

Data were analyzed using GraphPad Prism software version 10.6.1. Comparisons between the test and control groups were performed using an unpaired (independent) two-tailed t-test, assuming Gaussian distribution and equal variances. Data were entered as replicate values in separate columns for each group. Results are presented as mean ± standard deviation (SD), and statistical significance was defined as a p-value less than 0.05. Analyses were conducted using GraphPad Prism’s default unpaired t-test settings.

## Supporting information

**S1 Fig. Gating strategy for flow analysis.**

**S2 Fig. Flow analysis for cell surface expression of CD4, CXCR4 and CCR5 on HOS cells.**

**S3 Fig. Flow cytometry analysis of HIV 1 GFP reporter infection in CXCR4 expressing HOS cell lines.**

## Author contributions

**Conceptualization:** Beth Binnington, Donald R. Branch.

**Data curation:** Gizelle J. Lionel, Beth Binnington.

**Formal analysis:** Gizelle J. Lionel, Beth Binnington, Donald R. Branch.

**Investigation:** Gizelle J. Lionel, Beth Binnington.

**Methodology:** Gizelle J. Lionel, Beth Binninton, Jiazhen Jin, Alan Cochrane, Donald R. Branch.

**Project administration:** Beth Binnington, Donald R. Branch.

**Supervision:** Beth Binnington, Alan Cochrane, Donald R. Branch.

**Validation:** Gizelle J. Lionel, Beth Binnington.

**Visualization:** Gizelle J. Lionel.

**Writing – original draft:** Gizelle J. Lionel.

**Writing – review & editing:** Beth Binnington, Alan Cochrane, Donald R. Branch.

## Notes

### Competing Interest Statement

The authors have declared no competing interest.

