## Supplemental files for "A cell line model for the study of CD4-negative HIV-1 infection and latent virus reservoirs"

**Supplemental Materials**

**Detailed Methods**

**Supplemental information**

**Figure S1**

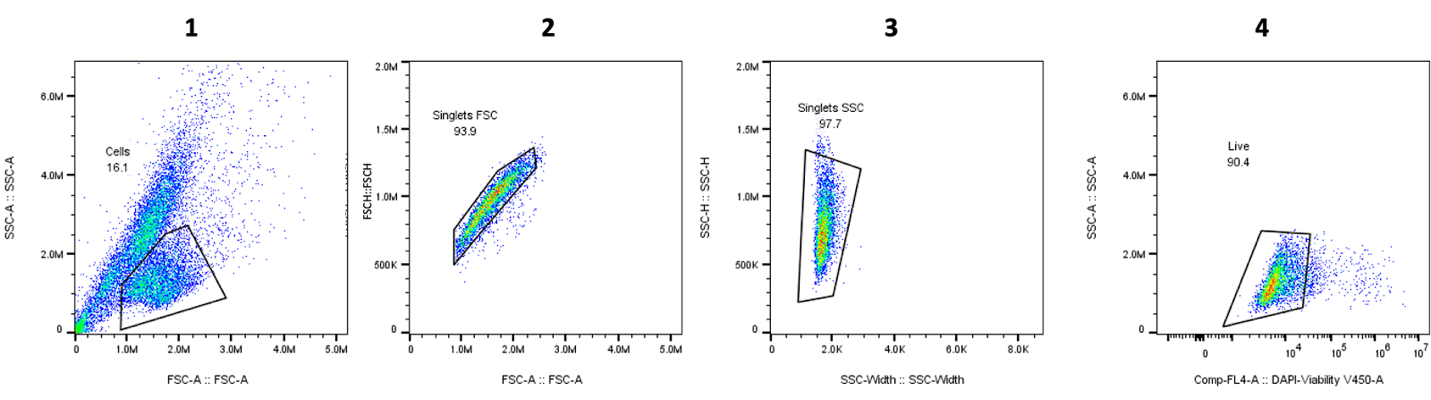

**Fig. S1.** Gating strategy for flow cytometry analysis. (1) SSC-A vs FSC-A, (2) FSC-H vs FSC-A, (3) SSC-H vs SSC-Width and (4) Viability gate.

**Figure S2**

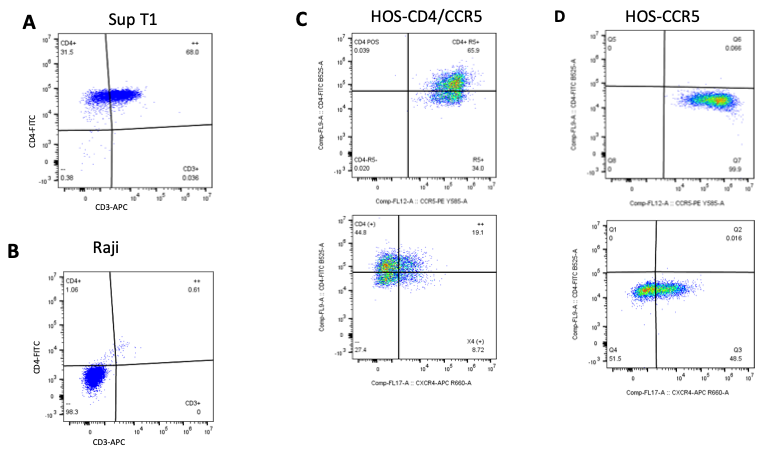

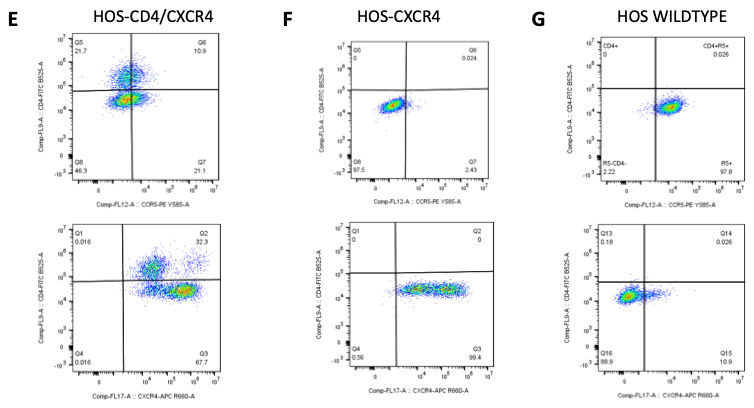

**Fig. S2.** **Flow cytometric characterization of CD4, CCR5, and CXCR4 surface expression in human osteosarcoma (HOS) cell lines, including SupT1 and Raji CD4‑staining controls.**
(A, B) Gating controls for anti‑CD4 staining: the CD4‑positive T cell line SupT1 (A, CD4 versus CD3) is 100% CD4‑positive, while the Raji B cell line (B, CD4 versus CD3) is CD4‑negative, confirming appropriate antibody specificity under the same flow conditions. (C–G) Representative dual‑parameter plots show surface expression of CD4 versus CCR5 (top row) and CD4 versus CXCR4 (bottom row) for HOS cell lines. Gates were set on live, single cells identified by forward‑ and side‑scatter and exclusion of doublets. HOS‑CD4/CCR5 (C) and HOS‑CD4/CXCR4 (E) cells express CD4 together with their respective coreceptor, with 66% of HOS‑CD4/CCR5 and 32.6% of HOS‑CD4/CXCR4 cells positive for CD4, while coreceptor expression approaches 100%. The CD4‑negative subpopulations in these lines are not attributable to inadequate anti‑CD4 staining, as demonstrated by the SupT1 and Raji controls (A, B). HOS‑CCR5 (D), HOS‑CXCR4 (F), and wild‑type HOS (G) cells are negative for CD4 but show high levels of CCR5 and/or CXCR4, consistent with endogenous CXCR4 expression reported as ubiquitous. Data were acquired from 5,000 events per sample and analyzed using FlowJo v10.9.0. These expression patterns confirm that CD4 is present only in CD4‑transduced HOS lines and support the use of HOS‑CD4+CCR5+ and HOS‑CD4+CXCR4+ cells for subsequent HIV entry and infection studies.

**Figure S3**

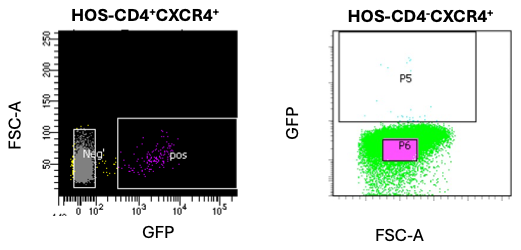

**Fig. S3.** **Flow cytometry analysis of HIV‑1 GFP‑reporter infection in CXCR4‑expressing HOS cell lines.** Representative flow cytometry plots show infection of HOS‑CD4^+^CXCR4^+^ (left) and HOS‑CD4^-^CXCR4^+^ (right) cells with an HIV‑1 GFP reporter -HXB2 env. Cells were infected at 5 × 10⁵ cells per T‑75 flask and analyzed by flow cytometry 70 h later. HOS‑CD4^+^CXCR4^+^ cells exhibited 2% GFP‑positive cells, and HOS‑CD4^-^CXCR4^+^ cells showed 0.1% GFP‑positive cells. GFP gating was set using uninfected control cells, and data were analyzed using standard flow cytometry software.
